# Assessing the Gene Regulatory Landscape in 1,188 Human Tumors

**DOI:** 10.1101/225441

**Authors:** C Calabrese, K Lehmann, L Urban, F Liu, S Erkek, NA Fonseca, A Kahles, H Kilpinen, J Markowski, PCAWG Group 3, SM Waszak, JO Korbel, Z Zhang, A Brazma, G Rätsch, RF Schwarz, O Stegle

## Abstract

Cancer is characterised by somatic genetic variation, but the effect of the majority of non-coding somatic variants and the interface with the germline genome are still unknown. We analysed the whole genome and RNA-Seq data from 1,188 human cancer patients as provided by the Pan-cancer Analysis of Whole Genomes (PCAWG) project to map *cis* expression quantitative trait loci of somatic and germline variation and to uncover the causes of allele-specific expression patterns in human cancers. The availability of the first large-scale dataset with both whole genome and gene expression data enabled us to uncover the effects of the non-coding variation on cancer. In addition to confirming known regulatory effects, we identified novel associations between somatic variation and expression dysregulation, in particular in distal regulatory elements. Finally, we uncovered links between somatic mutational signatures and gene expression changes, including *TERT* and *LMO2*, and we explained the inherited risk factors in APOBEC-related mutational processes. This work represents the first large-scale assessment of the effects of both germline and somatic genetic variation on gene expression in cancer and creates a valuable resource cataloguing these effects.

## Introduction

Cancer is characterised by extensive somatic genetic alterations. These variations can result in important cellular phenotypes with relevance for disease, including uncontrolled proliferation, immune evasion and metastasis (Weir, Zhao, and Meyerson 2004; Knudson 2002). Somatic mutagenesis is in part explained by environmental and intrinsic risk factors, with growing evidence for the relevance of the germline background of the patient (Pleasance et al. 2010; Jia, Pao, and Zhao 2014; Saini et al. 2016; Nik-Zainal et al. 2014). However, the interplay and the functional relevance of these different genetic factors are not sufficiently known.

One strategy to shed light on this question is association analysis with molecular readouts such as gene expression levels. Previous efforts using exome- and transcriptome sequencing data from The Cancer Genome Atlas (TCGA, Cancer Genome Atlas Research Network, Weinstein, et al. 2013) and the International Consortium of Cancer Genomes (ICGC, J. Zhang et al. 2011) have identified associations between somatic variants in coding regions and gene expression. Although these studies have helped to identify and characterise regulatory drivers, the role of the much larger number of non-coding somatic variants is not fully understood (Cancer Genome Atlas Research Network, Kandoth, et al. 2013; Kanchi et al. 2014). Recent studies have begun to address this by identifying genomic loci that are recurrently altered by somatic mutations, including their linkages to gene expression levels (Weinhold et al. 2014; Fredriksson et al. 2014; Smith et al. 2015; Ding et al. 2015). Additionally, substantial work has focused on the effects of variation in promoters of established cancer-genes, including *TERT* and *BCL2* (Weinhold et al. 2014; Fredriksson et al. 2014; Smith et al. 2015; Ding et al. 2015). However, thus far, a comprehensive analysis of associations between non-coding somatic variation and gene expression is missing.

We carried out joint genetic analyses that integrate coding and non-coding somatic variation with germline variants to investigate regulatory effects on gene expression levels in 27 cancer types. Building on 1,188 consistently processed genomes (whole genome sequencing, WGS) and transcriptomes (RNA-Seq) from the Pan-cancer Analyses of Whole Genomes (Campbell et al. 2017; PCAWG Consortium 2017) project, we have derived a detailed regulatory map that integrates different dimensions of germline and somatic variation, including single nucleotide variants (SNVs), somatic copy-number alterations (SCNAs) and signatures that capture differences in the prevalence of mutational processes across patients. Our approach combines complementary strategies, including allele-specific expression (ASE) analyses, somatic and germline expression quantitative trait loci (eQTL) mapping and the analysis of gene expression associations with global somatic signatures (**Fig. 1**).

**Fig. 1.**
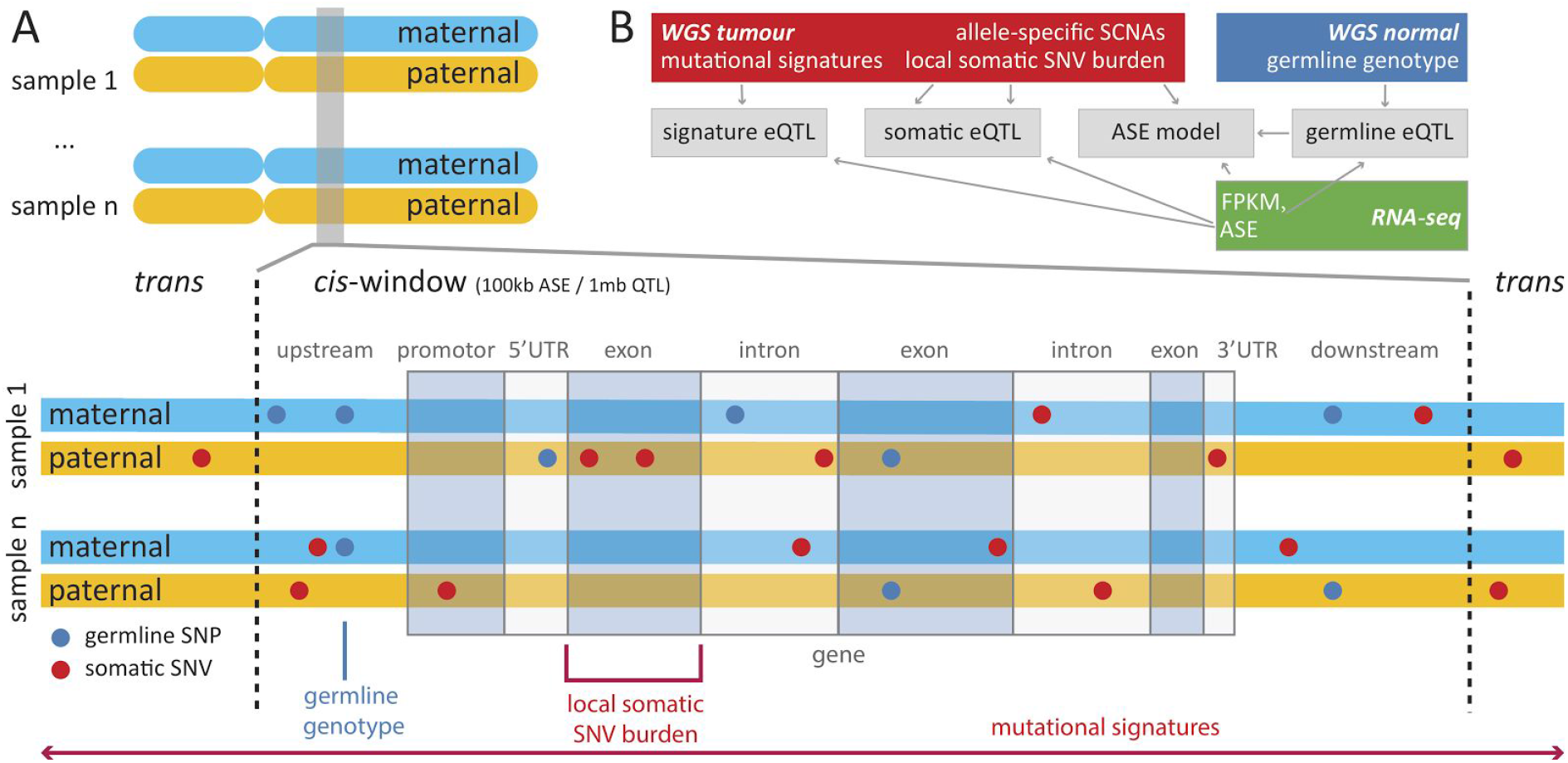
Integrative analysis approach for assessing the regulatory landscape in human cancers and overview of different classes of genetic variation considered here. **A)** *Cis* genetic effects of individual germline variants (blue) on gene expression were assessed using eQTL mapping. Somatic variants (red) were aggregated in mutational burden categories and assessed using i) ASE, ii) gene level *cis* eQTL and iii) associations with mutational signatures in patients. **B)** Data sources and processing steps, including germline and somatic variant calling from WGS data and allele-specific expression quantification from RNA-seq.

Collectively, our analyses provide a comprehensive picture of the regulatory landscape in human cancers, thereby identifying previously underappreciated associations between somatic regulation in distal regulatory elements and gene expression, as well as *de novo* functional annotations of mutational signatures. We report several associations that involve cancer-testis (CT) genes with known immunogenic properties (R.-F. Wang and Wang 2017; Scanlan et al. 2002), which exhibit high expression in sperm and some cancers but are repressed in healthy tissues (Simpson et al. 2005). Our results point at non-coding somatic dysregulation as a functional driver of carcinogenesis beyond mutations of the coding sequence and major contributor to inter- and intra-tumor heterogeneity.

## Results

We investigated 1,188 cases of 27 different tumor types obtained as part of the PCAWG working group (Campbell et al. 2017; Yung et al. 2017; PCAWG Consortium 2017), with WGS and RNA-Seq data (Yung et al. 2017; Whalley et al. 2017; Fonseca et al. 2017; PCAWG Group 1 2017; PCAWG Group 8 2017; PCAWG Group 3 2017, **Methods**). Across this panel, we quantified gene expression levels for 18,898 protein-coding genes (FPKM ≥ 0.1, in at least 1% of the patients, **Methods**), and allele-specific expression (**Methods**, **Fig. 1**).

### Germline Regulatory Variants in Cancer

We considered common germline variants (MAF≥1%) proximal to individual genes (± 100kb around the gene) to map eQTL across the entire cohort (**Fig. S2 A**). This pan-cancer analysis identified 3,509 genes with an eQTL (FDR ≤ 5%, hereafter denoted eGenes; **Methods, Table S2**), enriched in transcription start site (TSS) proximal regions as expected from eQTL studies in normal tissues (Bryois et al. 2014, **Fig. S2 B**). Analogous tissue-specific eQTL analyses in seven cancer types that were represented with at least patients identified between 106 (Breast-AdenoCA) and 472 eGenes (Kidney-RCC) (**Fig. S3 A, Table S2, Methods**).

To identify regulatory variants that are cancer-specific, we compared our eQTL set to eQTL maps from normal tissues obtained from the GTEx project (GTEx Consortium et al. 2017), adapting the strategy devised in Kilpinen et al. (2017). For each eQTL lead variant, we assessed the marginal replication in GTEx tissues (P<0.01, Bonferroni adjusted for 42 somatic tissues excluding cell lines, using proxy variants r^2^>0.8 for missing variants, **Methods**). For 87.5% of the eQTL that could be assessed in at least one GTEx tissue (2,982 of altogether 3,408 eQTL, **Fig. 2D**), this identified a replicating eQTL in at least one GTEx tissue, whereas 426 eQTL did not replicate in GTEx tissues, indicating cancer-specific regulation of gene expression (**Table S3**). One such example is *SLAMF9*, a member of the CD2 subfamily, with known roles in immune response and cancer (X. A. Zhang et al. 2003, **Fig. S4 A**). Similarly, we identified cancer-specific eQTL for genes with known roles in cancer such as *SLX1A*, an important regulator of genome stability that is involved in DNA repair and recombination and is associated with Fanconi Anemia (Saito et al. 2012; Medves et al. 2016, **Fig. S4 B**). The majority of these cancer-specific regulatory variants could not be explained by differences in gene expression level between cancer and normal tissues (328/426 exhibit at most a 2-fold increase in median gene expression compared to GTEx, e.g. *SLX1A*, **Fig. S3 B**), with a remaining set of 98 genes showing clear evidence of upregulation or ectopic expression (e.g. *SLAMF9*). This latter group contained immunoglobulin genes and nine CT genes including *RAD21L* (**Fig. 2, Fig. S3 B**). We also identified instances of eQTL that replicated in GTEx tissues but not in their corresponding normal tissues. One such example is *TEKT5*, which is expressed in our cohort but otherwise specific to testis in GTEx normal tissues, pointing to upregulation of selected genes in cancer (**Fig. S3 C-E**).

**Fig. 2.**
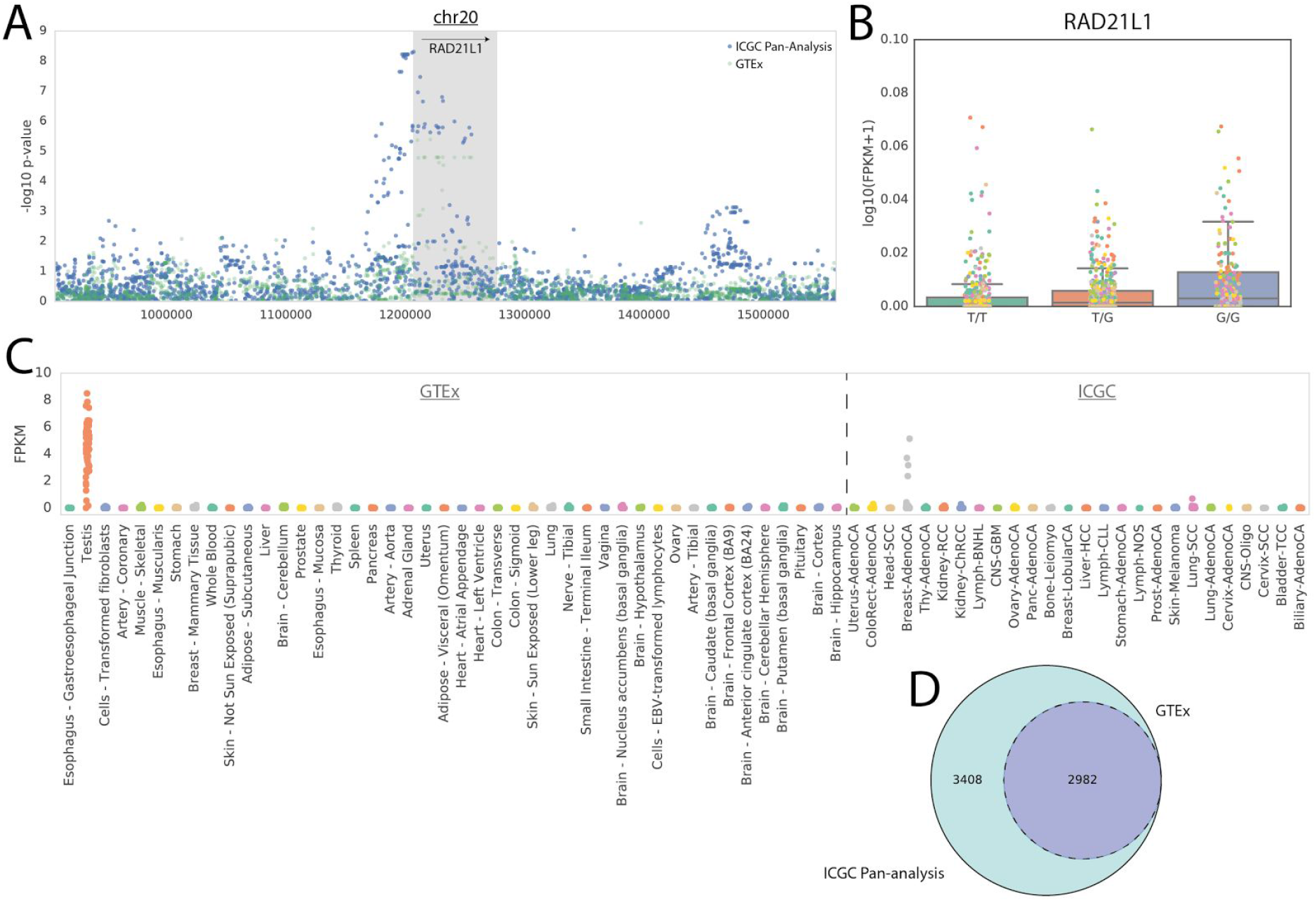
Germline eQTL analysis. **A)** Manhattan plot for *RAD21L1*, showing associations in the ICGC cohort (blue) and in GTEx testis tissue (green). **B)** Boxplots of the effect of the eQTL lead variant for *RAD21L* in the ICGC cohort. Colors denote cancer types of individual patients. **C)** Gene expression distribution of *RAD21L* across GTEx tissues and ICGC cancer types. *RAD21L* expression is increased in testis tissue, breast adenocarcinoma patients (‘Breast-AdenoCA’) and leukemia cell lines. **D)** The Venn diagram shows the number of eQTL identified in the ICGC cohort and the fraction of QTL that replicate in the GTEx cohort. 426 eQTL were specific to the ICGC cohort.

### Allele-specific Expression Captures Cancer-specific Dysregulation

To robustly identify genetic elements that contribute to somatic dysregulation of gene expression, we considered ASE, a locally controlled readout that enables assessing differential regulation between haplotypes in the same patient (Korir and Seoighe 2014). We quantified ASE at heterozygous germline variants, following Castel et al. (2015) (**Methods**). To maximise detection power, we aggregated ASE counts across heterozygous sites within genes, leveraging phased germline variant maps derived using a combination of statistical, copy-number (CN) and read-based variant phasing (**Methods**). This allowed quantification of ASE for between 588 and 7,728 genes per patient (median of 4,112 genes with at least 15 ASE reads in 1,120 patients, **Fig. S5**, **Methods**).

We tested for allelic expression imbalance (AEI) (FDR ≤ 5%, binomial test, **Methods**), finding substantial differences in the fraction of genes with AEI between cancer types (median percentages between 14.2% in Prost-AdenoCA and 46.8% in Lung-SCC among tumors; P=2.2x10^-13^ Mann-Whitney-Wilcoxon Test, **Fig. 3A**), and between AEI in cancer and the corresponding normal tissue (**Fig. 3B**). Cancers with extensive chromosomal rearrangements, including lung, breast and ovarian cancers, were associated with most frequent AEI events (**Fig. 3A, Fig. S6**), which is consistent with previous reports that have implicated SCNAs in allelic dysregulation in cancers (Ha et al. 2012).

**Fig. 3.**
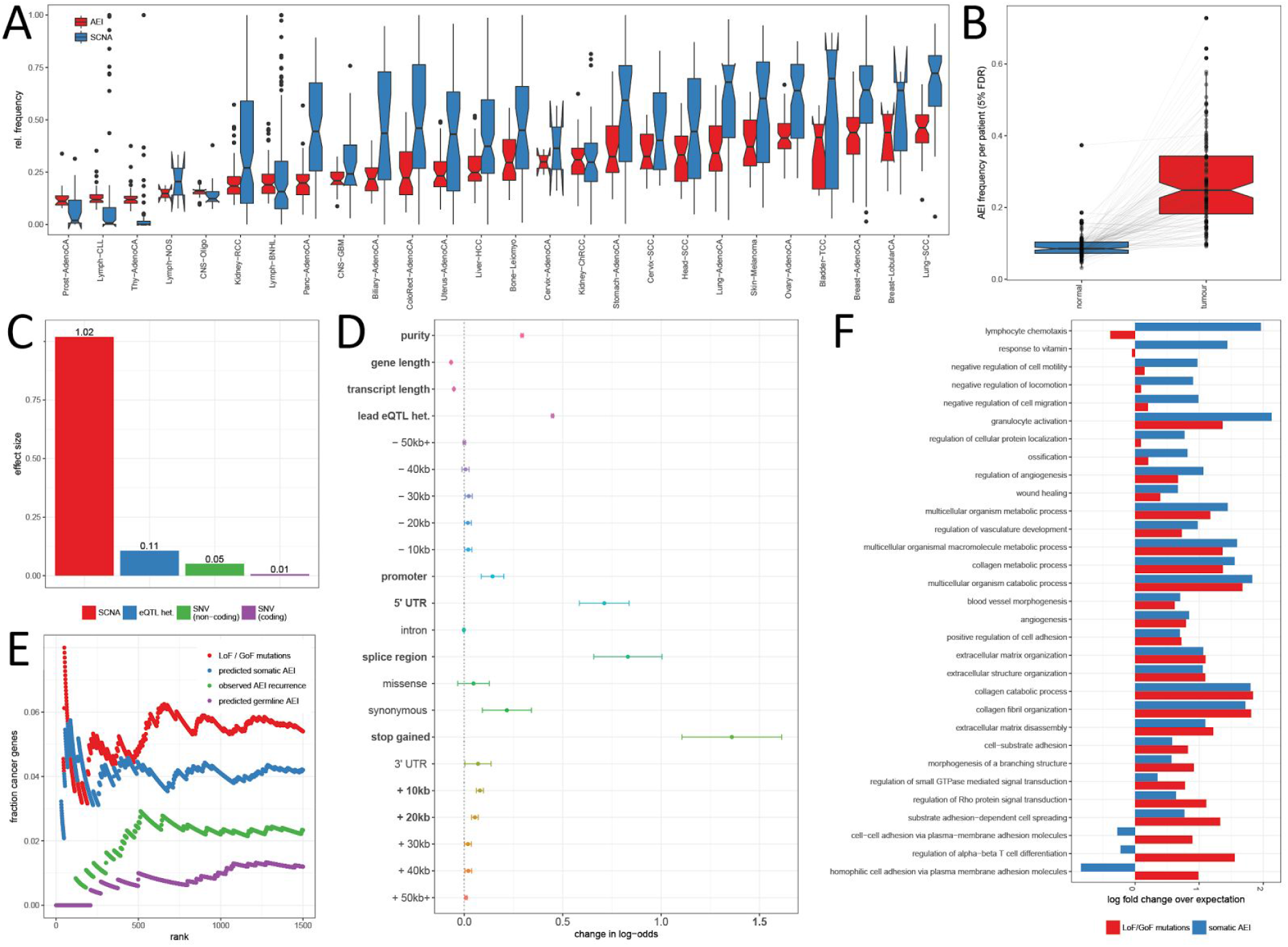
Allele-specific expression analysis. **A)** Distribution of the proportion of genes with significant allele-specific imbalance (AEI, FDR ≤ 5%) (red) and SCNAs (blue) across patients for different cancer types. Cancer types with prevalent chromosomal instability (frequent SCNAs) exhibit most frequent AEI. **B)** Proportion of genes with AEI (FDR ≤ 5%) in tumor and matched normal RNAseq patients across the cohort. **C)** Effect sizes of SCNAs, germline eQTL, coding and non-coding variants on AEI status. **D)** Relevance of individual somatic mutation types, germline eQTL and other co-variates for AEI status. Significant covariates (FDR ≤ 5%) highlighted in bold. **E)** COSMIC cancer gene enrichment for the four models of gene dysregulation: (i) average observed AEI frequencies (green); (ii) predicted germline AEI (purple); (iii) predicted somatic AEI excluding germline (blue), and (iv) LoF and GoF mutations (red). **F)** Enrichment of top 10% of genes ranked according to LoF/GoF mutations (red) or predicted somatic AEI (blue), using GO ontologies (FDR ≤ 5%).

Motivated by this, we used logistic regression to model the determinants of AEI (**Methods**), accounting for the presence or absence of germline eQTL, local allele-specific SCNAs and the mutational burden of proximal somatic SNVs, weighted by their respective cancer cell fraction and stratified into functional categories (upstream, downstream, promoter, 5’UTR, intron, synonymous, missense, stop gain, 3’UTR, **Fig. 1**, **Methods**). In aggregate, SCNAs accounted for 86.14% of the explained AEI variability, followed by germline eQTL (9.03%) and somatic SNVs (4.83%) (**Fig. 3C**). While cumulatively, non-coding variants were more relevant than coding variants (**Fig. 3C**), somatic protein truncating variants (‘stop-gained’) were the most predictive individually (**Fig. 3D**), which confirms the importance of nonsense-mediated decay (NMD) in cancer gene regulation (Lindeboom, Supek, and Lehner 2016). SNVs within splice regions, 5’ UTR and promoters were also strongly associated with AEI presence and we observed a global trend of decreased relevance of variants as a function of the distance from the TSS (**Fig. 3D**). We also considered a quantitative model on ASE ratios using phased variants as features, confirming downregulation of allelic expression by NMD (**Fig. S7 A-D**).

Using the trained model for AEI, we set out to characterise sets of genes with strong allelic dysregulation that can be attributed to different different genetic factors. We ranked genes according to average scores across the cohort, based on (i) the predicted AEI from the germline component and (ii) the predicted AEI from somatic components (SCNAs and SNVs) without germline effects. For comparison, we also considered (iii) the empirical AEI frequency in the cohort; and (iv) the burden of loss-of-function (LoF) and gain-of-function (GoF) mutations derived from genetic data only (**Fig. 3E**). When assessing these rank lists using known cancer genes (COSMIC, Forbes et al. 2017), we found cancer genes to be enriched among genes with high somatic AEI score (P≤0.005, Gene Set Enrichment Analysis, Subramanian et al. 2005, **Fig. S8 C**), however we observed no enrichment among genes with recurrent AEI (P=0.99, **Fig. S6 A**). As expected, genes with AEI due to germline eQTL were depleted for cancer genes (negative enrichment, P≤0.001, Fig. S8 B). Finally, consistent with the traditional definition of cancer genes based on recurrent mutations (Forbes et al. 2017), genes with recurrent LoF/GoF mutations were most strikingly enriched for the COSMIC census (P≤0.001, **Fig. S8 D**). The top 10% of genes of the somatic AEI scores were enriched for Gene Ontology (GO) categories with relevance to cancer, including chemotaxis, cell motility, locomotion and cell migration, which notably were absent when considering LoF/GoF mutations (**Fig. 3F**, Ashburner et al. 2000; Gene Ontology Consortium 2015a). These results suggest that somatic AEI could be used to prioritise regulatory variants to identify genes with roles in cancers, which extends previous observations in a single cancer (Ongen et al. 2014) to a pan-cancer setting.

Due to the strong effect of SCNAs on AEI we specifically investigated genes that were primarily dysregulated by SNVs. Of the 4,007 genes in the upper quartile based on the prevalence of overall AEI across all tumors, 1,843 genes exhibited SNV-linked AEI. When we ranked these genes based on the predicted AEI from SNVs, the top 10 genes were *FBXO5, ASPM, PSCA*, *CDKN1A*, *KIF20B*, *TP53* and *CLDN4*, which have previously been linked to cancer (Z. Wang et al. 2014; W.-Y. Wang et al. 2013; Gu et al. 2000; Abbas and Dutta 2009; Liu, Gong, and Huang 2013; Shang et al. 2012; Olivier, Hollstein, and Hainaut 2010), but also *EXO1*, *SYNE1* and *STON1-GTF2A1L*, genes that have not been prominently linked to cancer. *SYNE1* controls nuclear polarity and spindle orientation, which is upstream of NOTCH signaling in squamous lineage development (Lasorella, Benezra, and Iavarone 2014; Garraway and Lander 2013). Notably, based on the CADD analysis (Kircher et al. 2014), three melanoma cases preferentially expressed deleterious missense mutations (all CADD scores >25) in *SYNE1*, likely leading to a relative decrease of gene expression in these tumors. Based on the GEPIA web server (Tang et al. 2017), we also found that low *SYNE1* expression was associated with worse overall survival in TCGA melanoma patients (Log rank P=0.002, Hazard ratio=0.57, **Fig. S9 A**), providing further support for its relevance in disease. *EXO1* is known to be involved in mismatch repair and recombination, and exhibited significant AEI for a deleterious missense (CADD score 34, SIFT score 0, Kircher et al. 2014; Kumar, Henikoff, and Ng 2009) and a nonsense mutation in a colorectal adenocarcinoma patient. Similarly, TCGA colorectal adenocarcinoma patients with lower expression of *EXO1* showed worse overall survival (Log rank P=0.022, HR=0.57, **Fig. S9 B**), implicating *EXO1* as a potential tumor suppressor in human colorectal cancer. Consistent with this finding, *EXO1* knockout mice exhibited defects in DNA damage response and increased tumorigenesis (Schaetzlein et al. 2013).

Motivated by the observed cancer-specific germline regulation of CT genes, we investigated AEI in these genes. Notably, CT genes were depleted when considering the full somatic score (SCNAs and SNVs, 25/476 CT genes in the top 10% of genes, 48 expected, chi-square test, P=6x10^-4^), but enriched in the AEI score based on SNVs (66/476 CT genes in the top 10% of genes, 48 expected, chi-square test P=0.006). One potential explanation is that CT genes with low or no expression in differentiated tissues have to undergo somatic re-activation by SNVs before any subsequent SCNA can have an effect. To elucidate this, we used mutation timing data (Gerstung et al. 2017; PCAWG Group 11 et al. 2018, **Methods**), stratifying SNVs into the categories *early* (SNV occurred before SCNA at the same locus) and *late* (SNV occurred after SCNA at the same locus). We found strong over-representation of *early* SNVs in 329 out of 7525 CT gene-patient pairs (216 expected, chi-square test, P=3.96x10^-14^), suggesting that somatic re-activation of developmental genes through SNVs is selected for and precedes SCNA events at the same locus.

### Somatic eQTL Mapping Reveals Widespread Associations with Non-coding Variants

We next explored the effect of *cis* somatic variation on gene expression by aggregating somatic SNVs using local burdens in intronic and exonic genomic intervals (Gencode annotations, Harrow et al. 2012) and in consecutive genomic intervals within 1Mb from the gene boundaries (2kb regions, 1kb overlap, **Methods**), hereafter denoted as flanking regions.

Initially, we used these somatic burdens in conjunction with *cis* germline variants and SCNAs to decompose variation in gene expression of individual genes into their underlying components (**Methods, Fig. 4A**). Consistent with the ASE analysis, this identified SCNAs as the major source of variation (27.3% on average across all genes, **Fig. 4A**), followed by flanking somatic and germline variants. Notably, *cis* germline effects, although exhibiting smaller effects on average, were the largest variance component for 11,905 genes, compared to 3,568 genes which variation was primarily explained by somatic factors. Consistent with ASE analysis, we observed that non-coding somatic variants had more explanatory power than variants in coding regions.

**Fig. 4.**
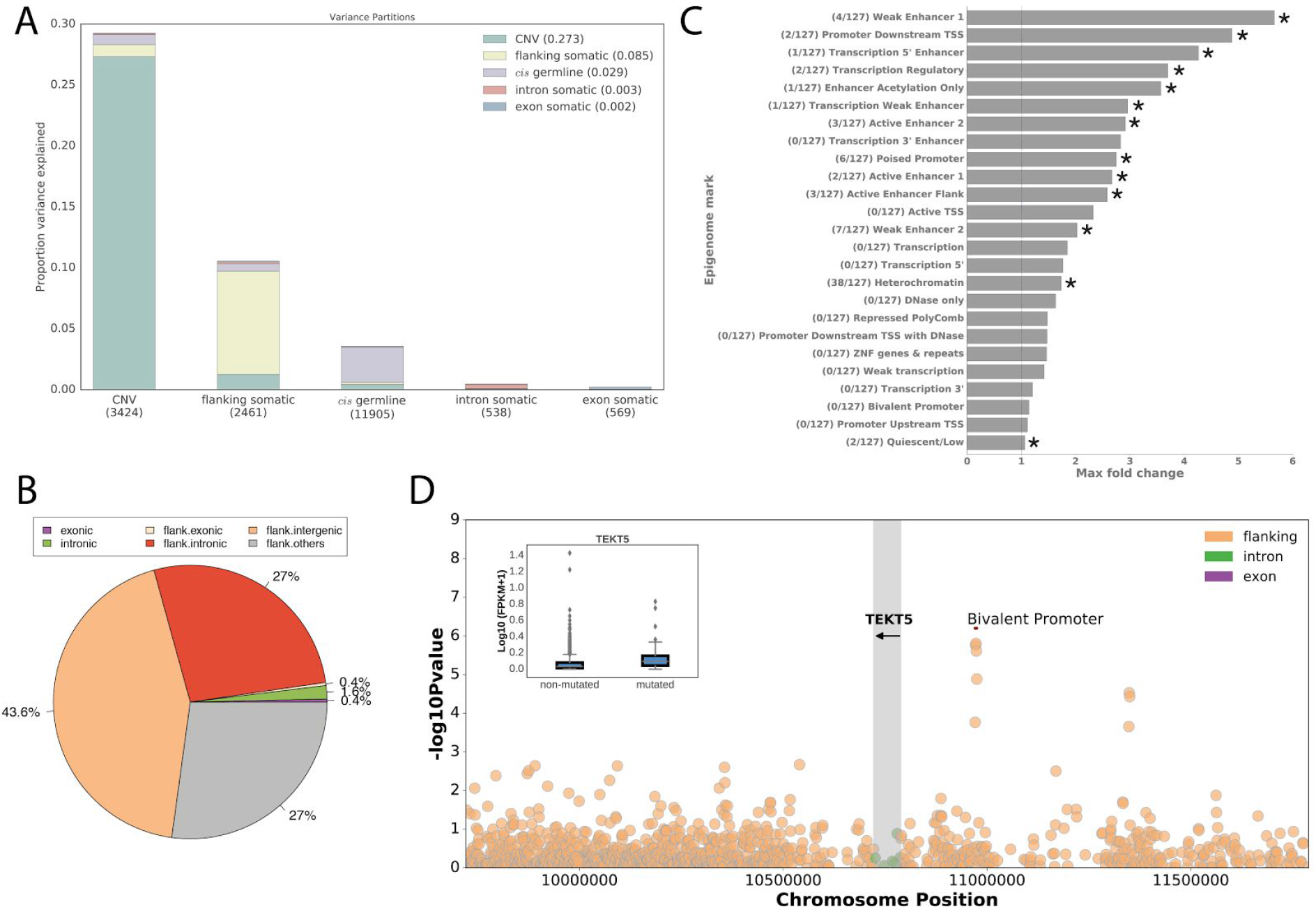
Somatic burden eQTL analysis. **A)** Variance component analysis for gene expression levels (**Methods**). Shown is the average proportion of variance explained by different germline and somatic factors for different sets of genes, considering genes for which the largest variance component are i) copy number effects (CNV), ii) somatic variants in flanking regions, iii) *cis* germline effects and iv) somatic intron and exon mutations, respectively. The number of genes in each set is indicated in parentheses. **B)** Breakdown of 567 genomic regions that underlie the observed *cis* somatic eQTL by variant category (Intronic = eGene intron; Exonic = eGene exon; Flank. = 2kb flanking region within 1Mb distance to the eGene start; Flank.intergenic = flanking region in a genomic location without gene annotations; Flank.intronic = flanking region overlapping an intron of a nearby gene; Flank.others = flanking region partially overlapping exonic and intronic annotations of a nearby gene). **C)** Maximum fold enrichment of epigenetic marks from the Roadmap Epigenomics Project across 127 cell lines. The number of cell lines with significant enrichments is indicated in parentheses (FDR ≤ 10%); asterisks denote significant enrichments in at least one cell line. **D)** Manhattan plot showing nominal p-values of association for *TEKT5* (highlighted in gray), considering flanking, intronic and exonic intervals. The leading somatic burden is associated with increased *TEKT5* expression (P=1.56x10^-06^; β =0.221) and overlaps an upstream bivalent promoter (red box; annotated in 81 Roadmap cell lines, including 8 ESC, 9 ES-derived and 5 iPSC cell lines). The inset boxplot shows a positive association between the mutation status and expression levels.

Next, we tested for associations between recurrently mutated intervals (burden frequency ≥ 1%) and expression levels of individual genes (18,708 protein coding genes; median 952 intervals per gene, **Fig. S10 C**), accounting for local and global differences in mutational burden across patients as well as tumor purity, cancer type, local SCNAs and other technical covariates (**Methods**). We assessed alternative strategies to estimate mutational burdens and found that weighted burden that accounts for variant clonality maximised detection power (**Fig. S11 A-D**). Genome-wide, this identified 649 somatic eQTL (FDR ≤ 5%; **Table S6**) in 567 genomic regions. Among these, 11 somatic eQTL were explained by the mutational burden in exons or introns, including genes with known roles in the pathogenesis of specific cancers such as *CDK12* in ovarian cancer (Bajrami et al. 2013; Ekumi et al. 2015), *PI4KA* in hepatocellular carcinoma (Ilboudo et al. 2014), *IRF4* in leukemia (Havelange et al. 2011), *AICDA* in skin melanoma (Nonaka et al. 2016), *C11orf73* in clear cell renal cancer (Bhalla et al. 2017) and *BCL2* and *SGK1* in lymphoma (Weinhold et al. 2014; Hartmann et al. 2016, Fig. S12 A-G). For the majority of 444 eGenes (70%), we observed associations with flanking non-coding regions (272 intergenic and 172 intronic regions). Hereby, 43.6% of the 567 unique genomic regions were intergenic and did not overlap any other annotated gene feature (**Fig. 4B**). The associated elements were generally mutated in two or more cancer types (Fig. S13, Table S6). Unlike germline eQTL, the associations tended to be located in distal regions (≥20kb, 88%) with on average larger effect sizes (| β |=3.3) than associations in regions proximal to the TSS (| β |=1.4) (**Fig. S14**), which points to the relevance of somatic mutations at distal regulatory elements.

Motivated by this, we tested lead flanking regions for enrichments in cell type specific regulatory annotations in 127 cell types from the Roadmap Epigenomics Project (Roadmap Epigenomics Consortium et al. 2015) as well as transcription factor binding sites (TFBS) from ENCODE (8 cancer cells and one embryonic stem cell line, ENCODE Project Consortium 2012), comparing the overlap of true associations to distance and burden frequency matched random regions (**Methods**). This identified a significant enrichment (FDR ≤ 10%) of 13 out of 25 epigenetic annotations (**Fig. 4C**, **Table S7**), including poised promoters, weak and active enhancers and heterochromatin in more than two cell lines (**Fig. 4C**), but no significant enrichment of TFBS (**Tables S8**).

Poised or bivalent promoters are a hallmark of developmental genes and prepare stem cells for somatic differentiation (Lesch and Page 2014). Re-activation of poised promoters is one mechanism of upregulation of developmental genes in cancer, including CT genes (Bernhart et al. 2016). CT genes were marginally more frequent among genes with somatic eQTL than expected (45/982, P=0.06, Fisher’s exact test), and we observed an enrichment for eQTL in bivalent promoters for the CT genes (P=0.04, Fisher’s exact test). One such gene is *TEKT5*, an integral component of sperm, that has been found to be aberrantly expressed in a variety of cancers (Hanafusa et al. 2012). We observed a positive association between *TEKT5* expression and somatic mutational burden (prevalently observed in non-Hodgkin lymphoma patients) in a bivalent promoter site close to the 5’ end of the gene (**Fig. 4D**). The prevalence of developmental genes among the somatic eGenes was also consistent with a global enrichment (FDR ≤ 10%) for GO categories related to cell differentiation and developmental processes (**Table S9**). Together with the ASE analysis, these results emphasise the relevance of non-coding genetic variation for changes in gene expression and the importance of re-activation of developmental genes through somatic mutations for cancer progression.

### Associations between global somatic mutational signatures and gene expression

In addition to associations with *cis* elements proximal to genes, it is plausible that global differences in mutational states between patients are associated with cell phenotypes, including gene expression levels. Global variations in mutational patterns can be quantified using mutational signatures which tag mutational processes specific to their tissue-of-origin and environmental exposure (Alexandrov et al. 2013). However, the relationship between mutational signatures and gene expression levels is unknown.

We considered 28 mutational signature readouts (Forbes et al. 2017) derived using non-negative matrix factorization of context-specific mutation frequencies in the PCAWG cohort (PCAWG-7 beta 2 release, PCAWG Group 7 et al. 2018). We tested for associations between patient-specific signature prevalence and expression levels of individual genes, accounting for total mutational burden and additional technical as well as biological confounders (18,831 genes, FPKM filtering of genes across 1,159 patients, **Methods**). Across all signatures, this identified 1,176 genes associated with at least one signature (FDR ≤ 10%, **Fig. S15**, **Methods**), a markedly different set of genes compared to genes associated with the total mutational burden that ignores the assignment to different signatures (**Table S10 F**). Lymphoma Signature 9 was associated with the largest set of genes, followed by the smoking-related Signature 4 (**Fig. 5A, Tables S10 A,E, Fig. S16 D**).

**Fig. 5.**
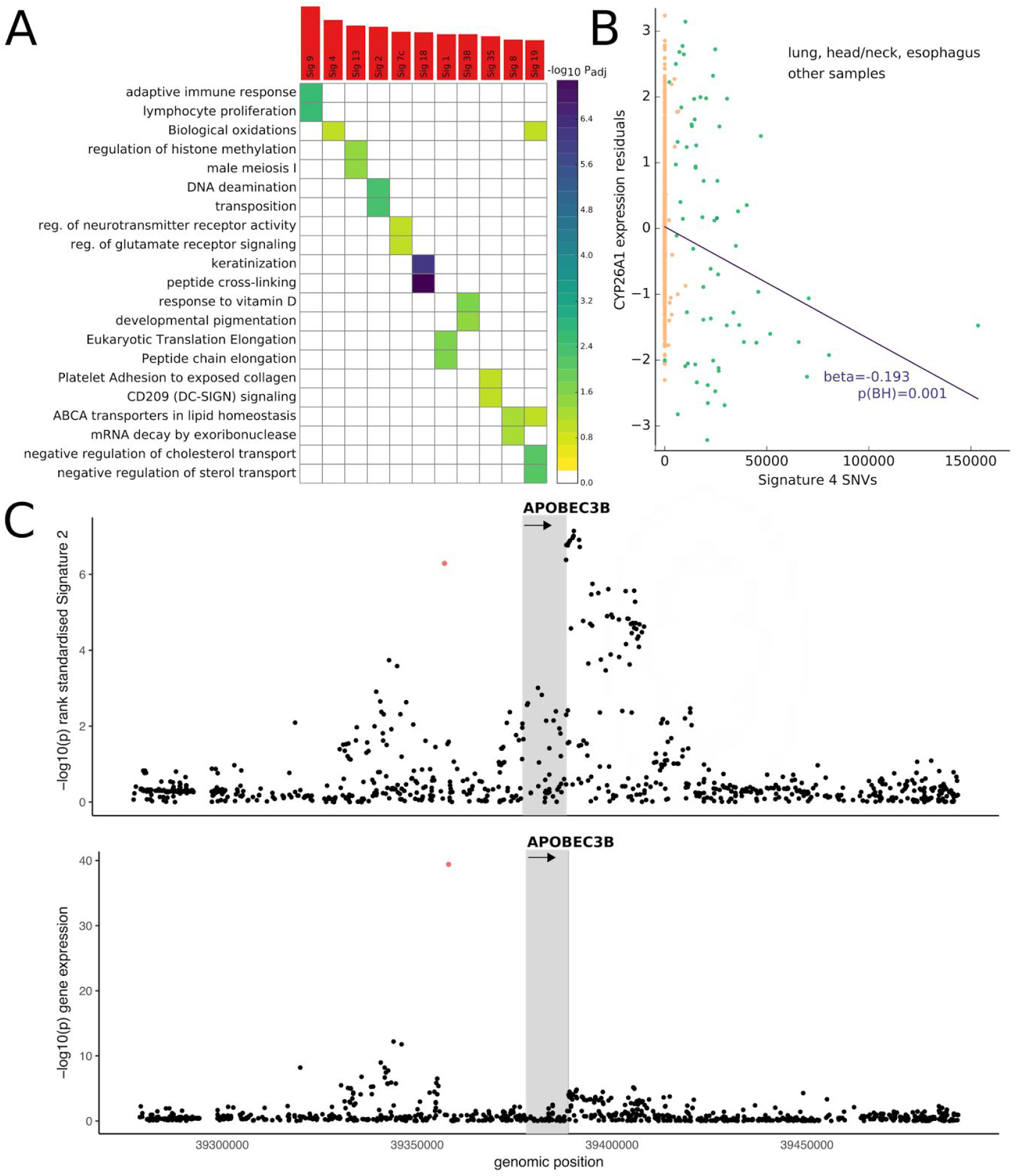
Associations between mutational signatures and gene expression. **A)** Summary of significant associations. Top panel: Total number of associated genes per signature (FDR ≤ 10%). Bottom panel: Enriched GO categories or Reactome pathways for genes associated with each signature (FDR ≤ 10%, significance level encoded in color, −log_10_ P_adj_). **B)** Representative signature-gene association, depicting a negative association between *CYP26A1* expression and Signature 4. **C)** Manhattan plots of associations between *cis* germline variants proximal to *APOBEC3B* (plus or minus 100kb from the gene boundaries) and Signature 2 (top panel) or *APOBEC3B* gene expression level (bottom panel). The gray region denotes the gene body, the orange variant the lead eQTL variant rs12628403.

While aetiologies for some mutational signatures are well understood, others have not previously been characterised. To annotate the signatures, we considered 18 signatures with at least 20 associated genes and tested for enrichments using GO and Reactome Pathways (Fabregat et al. 2016; Milacic et al. 2012) categories. Of these, 11 signatures were enriched for at least one category (FDR ≤ 10%, **Table S10 D**), revealing several associations that were consistent with known aetiologies (**Fig. 5A**). For example, Signature 9 is known to be active in certain lymphomas and leukemias (Forbes et al. 2017), and was associated with 354 genes enriched for lymphocyte/leukocyte-related processes and immune response, including *TCL1A*, *LMO2* and *TERT* (P=1.22x10^-10^, 6.84x10^-10^, and 1.96x10^-09^). Smoking Signature 4 (Forbes et al. 2017) was associated with 119 genes, enriched for biological oxidation associated processes (e.g. processing of benzo[a]pyrene) and including the gene *CYP24A1*, which is known to be down-regulated in tobacco-smoke exposed tissue (**Fig. 5B**, Woenckhaus et al. 2006). We further identified 70 genes associated with the APOBEC Signature 2 (Forbes et al. 2017), which were significantly enriched for DNA deaminase pathways.

Among those signatures with previously unknown aetiology, our results link Signature 8, which is primarily prevalent in medulloblastoma (Forbes et al. 2017), to a set of 25 genes enriched for ABCA-transporter pathways, which are targeted by drugs that are in clinical trials for medulloblastoma (Milacic et al. 2012; Ingram et al. 2013). Similarly, Signature 38, which is correlated with the canonical UV Signatures 7 (e.g. 7a: r^2^=0.375, P=5x10^-40^, Fig. S16 C), was linked to melanin processes (**Fig. 5A**). Melanin synthesis is known to subject melanocytes to oxidative stress by involving main oxidation reactions and superoxide anion/hydrogen peroxide generation (Kvam and Tyrrell 1999; Denat et al. 2014). Our data linked Signature 38 to the oxidative stress promoting gene Tyrosinase (*TYR*, P=1x10^-4^, Jimbow et al. 2001). As Signature 38 is characterised by an excess of C>A mutations, a typical product of reactive oxygen species (ROS) mediated by activity of 8-hydroxy-2’-deoxyguanosine (DFG, 2010), it might represent DNA damage indirectly caused by UV after direct sun exposure due to oxidative damage (Premi and Brash 2016), with *TYR* as a possible damage mediator.

The cause-and-effect relationship of correlated somatic variations and gene expression changes are not clear *a priori*. To begin addressing the directionality of these linkages, we explored the utility of germline variation as an anchor to gain directed mechanistic insights. Building on our eQTL map, we queried germline eQTL lead variants of mutational signature-associated genes and tested for associations between these variants and the corresponding mutational signature. This eQTL-guided approach entails substantially fewer tests than genome-wide analyses that consider all germline variants (Waszak et al. 2017; PCAWG Group 8 2017). Among 1,176 signature-linked genes, 197 also had a germline eQTL, but only eQTL variant rs12628403 was associated with the respective signature (FDR ≤ 10%, multiple testing over 197 association tests, **Table S10 E**). This germline variant is a known germline predisposition factor for Signature 2 prevalence (Middlebrooks et al. 2016), an eQTL for *APOBEC3A/B* and significantly associated with Signature 2 prevalence in our cohort (P=5.13x10^-7^, **Fig. 5C**). Colocalisation analysis (Wallace 2013) confirmed that the variant rs12628403 is a plausible genetic determinant of both, expression levels of *APOBEC3A/B* and Signature 2 (**Table S10 E**, **Methods**). Finally, we carried out mediation analysis (Baron and Kenny 1986; Preacher and Hayes 2004) to formally test whether expression levels of *APOBEC3A/B* confer the genetic effect to the signature (**Methods**). We found that *APOBEC3B*, *POTEI* (both P<10^-10^) and *APOBEC3A* (P=0.004) expression levels conferred the effect of rs12628403 to Signature 2. The proportion of the effect of the variant on the signature that is mediated by only *APOBEC3B* expression is a remarkable 87.11% (nonparametric bootstrapping, 1,000 simulations, **Fig. S17**, **Methods**).

## Discussion

The regulatory landscape of cancer is highly heterogeneous, cancer type specific and influenced by the germline background. This study provides a comprehensive picture of how different germline and somatic variations alter gene expression levels. Our results show that coregulation of the same genes by multiple different types of variants is common in cancer (**Fig. S1**). Here, we have assessed the relative magnitudes of these effects (**Fig. 4A**). Previous studies have been limited by the lack of whole genome sequencing data, which is essential for identifying contributions of non-coding variants to gene expression variability. Indeed, our analysis which is based on data from the currently largest cohort of matched tumor WGS and RNA-Seq data of 1,188 patients, demonstrates that the impact of non-coding variation can be profound.

We have produced comprehensive across-tissue and tissue-specific germline eQTL maps and have identified associated genetic risk variants. By comparing these cancer eQTL maps to the GTEx catalogue, we have observed substantial overlap between cancer and normal tissues, but also a smaller number (12.14%) of potentially interesting cancer-specific eQTL. This estimate is conservative since we have assessed replication in any GTEx tissues and not only in the corresponding normal tissues. In selected cases we have observed that eQTL in cancer mimic regulatory effects in other non-corresponding tissues, in particular testis-specific genes (**Fig. 2A**).

In parallel to germline eQTL, we have considered the effect of somatic mutational burden in different genomic regions on gene expression changes by building a systematic map of somatic eQTL. Our analysis accounts for variation in clonality as well as local hypermutations, thereby identifying likely causal associations between somatic burden and gene expression. However, this approach has limitations and we cannot rule out that a fraction of the associations we have identified are due to technical or biological factors that jointly affect gene expression and local mutation rates. We also note that an analysis of cancer type specific somatic eQTL is currently not feasible with the given sample size.

We have used ASE readouts for integrated modelling of genetic variation in *cis* and fine-grained characterisation of the genetic elements that have the largest regulatory effects. We have demonstrated the extent to which AEI follows allelic imbalance on the genomic level. While ASE is sensitive to heterozygous genetic variation, the considered phased somatic mutation set is based on read phasing, which has only been possible for around 20% of all SNVs. Further, ASE readouts can only be derived in cases with at least one heterozygous germline variant in the gene in question, reducing overall sample size and hindering gene-level associations.

Our analysis suggests somatic re-activation of CT genes through SNVs, supported by mutation timing analysis. CT genes were also enriched in cancer-specific germline and somatic eGenes linked to SNV burden in nearby bivalent promoters. Somatic eQTL were further enriched for tissue development and differentiation. Due to the low number of expressed CT genes these findings need to be evaluated on additional data. However, we have found CT gene implicated in three out of four independent analyses (germline eQTL, somatic eQTL and ASE), reinforcing the potential impact of these findings.

Mutational signatures capture global variations across individuals, for example due to environmental factors or exogenous damage, which are distinct from local somatic variations assessed using QTL mapping and ASE. We have explored the utility of associations between these mutational signatures and gene expression levels, thereby deriving *de novo* annotations of signatures with previously unknown roles. Finally, we have carried out proof of concept analyses to integrate germline factors, somatic variants and gene expression, thereby unpicking the molecular chain of events for the common APOBEC mutational process and its germline component. Due to the tissue specificity of mutational signatures, it will be important to conduct similar analyses for individual cancer types and in cohorts with larger sample sizes.

This is the first large-scale study assessing the effects of both germline and somatic genetic variation on gene expression from WGS data in a pan-cancer setting. The somatic and germline eQTL resources will be a valuable resource to address a wide range of downstream analyses, providing a comprehensive overview of gene expression determinants in cancer and insights into the underlying biology. The systematic assessment of regulatory non-coding genetic variation significantly improves our understanding of the aetiology and functional implications of intra- and inter-tumor heterogeneity and the selective forces applied to these heterogeneous genomes.

## Acknowledgements

L.U., R.F.S. and O.S. received support from core funding of the European Molecular Biology Laboratory and the European Union’s Horizon2020 research and innovation programme (grant agreement number N635290). K.L., A.K. and G.R. received core funding from Memorial Sloan Kettering Cancer Center (New York) and from ETH Zurich. R.F.S. and J.M. received support from the Helmholtz Foundation and the Max Delbrueck Center for Molecular Medicine. F.L. and Z.Z. received support from National Natural Science Foundation of China (grant agreement numbers 31530036 and 81573022). C.C, N.F. and A.B. received support from core funding of the European Molecular Biology Laboratory and from the EU FP7 Programme projects EurocanPlatform (grant agreement number 260791) and CAGEKID (grant agreement number 241669).

## Contributions

The authors are listed according to their contributions. Senior contributors are listed in parentheses and in alphabetical order.

Data Preprocessing: Lehmann K, Calabrese C, Kahles A, Fonseca N, (Brazma A, Rätsch G, Stegle O)

Germline eQTL mapping/analysis: Lehmann K, Calabrese C, (Rätsch G, Stegle O)

Somatic eQTL mapping/analysis: Calabrese C, Lehmann K, Fonseca N, Urban L, (Brazma A, Rätsch G, Schwarz RF, Stegle O)

ASE preprocessing: Schwarz RF, Erkek S, Waszak S, (Schwarz RF, Korbel J, Stegle O, Zhang Z)

ASE analysis: Schwarz RF, Urban L, Liu F, Kilpinen H, Markowski J, (Schwarz RF, Stegle O, Zhang Z)

Signature analysis: Urban L, (Schwarz RF, Stegle O)

Germline/Signature interface: Urban L, Lehmann K, (Schwarz RF, Stegle O)

